# Benchmarking Boltz-2 for Screening of Therapeutic Antibody-Antigen Interactions

**DOI:** 10.64898/2026.05.13.724924

**Authors:** Alexandra Fieux-Castagnet, Julian Waton, Alina Glukhonemykh, Eric Snow, Roshini Ashokkumar, Jess Fleming, David Champagne, Thomas Devenyns, Alex Peluffo, Chris Anagnostopoulos

## Abstract

Protein structure prediction models (such as AlphaFold, Chai, Boltz) have transformed structural biology and are increasingly explored for drug discovery; however, their utility for large-scale screening of antibody-antigen (AB-AG) interactions remains unclear, particularly for distinguishing true binding from non-binding pairs at scale. To our knowledge, there has not been an exhaustive exploration of Boltz-2 inference settings on this high impact problem, and in this paper we set out to describe and implement a novel benchmarking framework that can accelerate progress in the field. We evaluated Boltz-2 (NVIDIA NIM implementation) on 519 therapeutic monoclonal antibodies from Thera-SAbDab, pairing each antibody with its cognate target and a randomly assigned non-cognate antigen. We developed a novel evaluation framework that systematically captures variability across stochastic seeds while benchmarking different inference settings, including datasets with and without crystallographically resolved antibody structures. Across settings, Boltz-2-derived confidence metrics showed weak, though above-chance, discrimination (0.5 < ROC-AUC < 0.60). Among evaluated metrics, the minimum value of the interface predicted TM-score (ipTM-min) across seed-samples, captured the strongest signal. Interestingly, additional feature aggregation and multivariate modelling provided little to no improvement. Increasing the number of stochastic predictions yielded front-loaded gains, with diminishing returns beyond ∼15–20 seed-samples, suggesting limited value of extensive sampling in practical workflows. Notably, inference without multiple sequence alignments (MSAs) slightly improved performance on non-crystallized antibodies (ΔAUROC ≈ +0.027) while reducing runtime by ∼8 seconds per prediction compared to shallow MSA settings. Overall, these results indicate that off-the-shelf confidence metrics from general-purpose structure prediction models may be insufficient for reliable target-antibody screening and highlight the need for task-specific optimization, while confirming that modest amounts of sampling can be helpful, but not in itself sufficient to improve performance significantly as gains plateau relatively quickly.

## Introduction

Recent advances in deep learning–based protein structure prediction have transformed structural biology and are increasingly influencing modern drug discovery workflows. Methods such as AlphaFold and related AI-driven modelling approaches enable the prediction of protein structures and complexes directly from their sequence, supporting applications ranging from protein-protein interaction and structure-based drug design to biologics engineering and target identification.^1–4^ These approaches have substantially accelerated structural analysis and reduced dependence on experimental structure determination, enabling rapid computational exploration of biomolecular interactions relevant to therapeutic development.

Therapeutic antibodies represent a particularly important and challenging test case for evaluating the practical utility of computational interaction prediction. Antibodies account for a substantial proportion of approved biologic therapeutics and play a central role in modern precision medicine strategies.^5^ Their discovery and optimization require the screening of large combinatorial libraries of candidate molecules while minimizing off-target interactions, nonspecific binding, and polyspecificity that could compromise safety or efficacy.^6–8^ As a result, computational methods capable of identifying selective antibody-antigen (AB-AG) interactions prior to experimental validation could significantly accelerate early-stage antibody discovery and improve candidate prioritization, complementing active learning frameworks.

Recent diffusion-based multimer models, AlphaFold2/3, Chai1/2, Protenix and Boltz-1/2, offer architectural advances that enable direct multi-chain complex modelling without separate docking steps.^9–13^ Boltz-2 provides an open-source, permissive, scalable implementation optimized for efficient inference and produces both predicted complex structures and a suite of confidence metrics, including interface predicted TM-score (ipTM), predicted TM-score (pTM), per-residue predicted local distance difference test (pLDDT), and predicted distance error (PDE). These metrics are widely used as proxies for structural plausibility and interface quality and are commonly used to assess predicted protein complexes.^14^ At the time of this study, Boltz-2 represents one of the highest-performing open-source models, making it a practical candidate for deployment in real-world antibody discovery workflows.^13^

Accurate structural prediction alone does not imply functional binding, and screening applications require reliable discrimination between binders and non-binders. Note that the focus of this work is on-target binding and not off-target binding, i.e. we are classifying true binders from similar non-binders from the same distribution of targets. Identifying optimal inference settings for structure-based antibody screening requires consideration of several key parameters that jointly influence predictive performance. These include (i) the interpretation of model-derived confidence metrics for binding discrimination, (ii) the use of evolutionary information such as multiple sequence alignments (MSAs), and (iii) the extent of stochastic sampling during model inference. Together, these factors define a multidimensional optimization space in which improvements in structural prediction accuracy (which is the primary goal of these models) do not necessarily translate into improved discrimination between binders and non-binders, motivating systematic evaluation of their individual and combined effects.

In practice, heuristic thresholds on confidence metrics— such as ipTM values of approximately 0.7—are often used to infer likely binding interactions.^15,16^ However, the validity of such thresholds for distinguishing true AB-AG binders from non-binders in realistic screening settings has not been systematically evaluated. Recent evidence in nanobody systems suggests that structurally plausible AB-AG complexes predicted by AI-based methods may not necessarily correspond to biologically meaningful binding interactions, underscoring the need for task-specific validation of these models in realistic discovery settings.^17^ Emerging observations further suggest that commonly used confidence metrics may exhibit weak discriminative power in this setting, motivating systematic evaluation.

The role of evolutionary information introduces further complexity. MSAs provide coevolutionary constraints that significantly enhance monomer and multimer structure prediction accuracy in AlphaFold2.^18–20^ In particular, AB-AG interactions often lack strong inter-chain co-evolutionary signal, as antibodies are artificially selected binders rather than co-evolved partners.^21–23^ This limitation may reduce the reliability of traditional structure-prediction confidence metrics when applied to antibody discovery problems. Moreover, the practical value of MSA usage in screening settings remains uncertain, particularly given trade-offs between computational cost and predictive performance. Consistent with this limitation, recent benchmarking on unseen complexes indicates that AlphaFold2 exhibits diminished accuracy in this context, with improvement driven primarily by increased sampling rather than by intrinsic co-evolutionary signal.^24^

In addition to MSA usage, stochastic inference and sampling variability represent important practical considerations in modern diffusion-based structure prediction models. Multiple independent predictions generated from different random seeds may produce a distribution of predicted structures with varying interface quality. AlphaFold3 has shown enhanced interface recovery under expanded sampling regimes (up to 1000 seeds).^9^ However, in the context of large-scale screening, where thousands of candidate interactions must be evaluated, increased sampling introduces a trade-off between potential gains in discriminative performance and substantially higher computational cost, and it remains unclear whether these gains are sufficient to justify the added expense. In exploring the practical utility of structure-based screening, it is therefore important to assess whether gains from additional sampling are substantial or exhibit diminishing returns beyond a limited number of predictions. Given prior observations of limited discriminative performance under fixed inference settings, it is important to assess whether gains from additional sampling are substantial or, instead, exhibit diminishing returns beyond a limited number of predictions.

In this study, we evaluate the structural confidence metrics outputted by Boltz-2, chosen at the time as the best permissive open-source model, for their ability to distinguish experimentally validated AB-AG binding pairs from randomly generated non-binding interactions in an insilico screening environment. Using predicted complex structures and associated confidence scores, we assess discrimination performance using receiver operating characteristic (ROC) analysis and area-under-the-curve (AUC) metrics. We further examine how inference choices, including sampling depth, model-derived confidence metrics (e.g., ipTM, pTM), MSA usage, and dataset composition, impact performance under practical screening constraints. Ultimately, this study provides a systematic evaluation of the practical utility of Boltz-2 for AB-AG screening and identifies key limitations that must be addressed before such models can support large-scale discovery workflows.

This article is organized into the following sections: Experimental Design, Experimental Results, Discussion, and Methods. Together, these sections provide an outline of the research design, present detailed analyses of the findings, interpret how the results contribute to the broader field, and describe the methodologies used.

## Experimental Design

This study evaluated the effects of inference regime and aggregation-depth parameter configurations in Boltz-2 on AB– AG binding prediction, using a controlled validation framework with known ground-truth on-target binding relationships. A curated dataset of 519 monoclonal antibodies from Thera-SabDab^25^ and 225 unique protein targets from UniProt was assembled.^26^ Each antibody was modelled with its cognate antigen and with one randomly reassigned non-cognate antigen drawn from the same target pool with replacement. By sampling negative controls from the identical distribution of targets and controlling for length and class, this design enables isolation of native vs negative control discrimination rather than dataset-driven bias. This section has a high-level description of the experimental approach; for a more detailed presentation, see the Methods section.

### Structure generation and parameter regimes

Complex structures were predicted using the NVIDIA Inference Microservices (NIM) implementation of Boltz-2. For each AB–AG pair, we generated 40 predictions by running 8 random seeds with 5 diffusion samples per seed (8 × 5). Two inference regimes were evaluated: (i) no-MSA and 32-MSA, chosen because MSAs have proven to be valuable in structure prediction but reduced to length 32 due to computational reasons.^27^

### Scoring and feature configurations

For each generated prediction, we extracted Boltz-2 output metrics (confidence score, ipTM, etc.) and constructed features by aggregating across the available predictions for that pair over samples and seeds. We evaluated two feature configurations: (1) individual metrics (e.g. ipTM) and (2) a full feature set derived from Boltz-2 confidence outputs.

### Discrimination task and evaluation

Discrimination performance was evaluated by distinguishing native pairs from negative controls using two complementary approaches. First, we assessed the ability of aggregated model confidence metrics (e.g., minimum ipTM), computed across multiple seed–sample predictions (8 seeds × 5 samples, treated as stochastic draws), to discriminate between classes via ROC–AUC. We explored several aggregation strategies to summarize these replicate predictions (e.g. minimum, maximum, etc.).

Second, we trained supervised classifiers on a set of aggregated features (including ipTM, confidence scores, and other outputs) and evaluated their performance using ROC– AUC across repeated train/test splits. A label-permutation control was included to establish a chance-level baseline.

All analyses were conducted on both the full dataset and a structure-excluded subset (excluding antibodies with crystallographic structures) to assess robustness to potential overlap with known structures (since Boltz-2 was trained on crystallographic structures released before June 2023).^13^

### Sampling-depth sensitivity analysis

To quantify sensitivity to diffusion-sampling depth, we treated the 40 predictions per pair as a fixed pool and evaluated performance as a function of the number of aggregated predictions. For each sampling depth, we subsampled seed–sample predictions, recomputed aggregated features, and repeated the classification evaluation. This produced a convergence profile of discrimination performance as sampling depth increased. This is described in and further explored in the Methods section.

## Experimental Results

### Baseline estimator performance

Out of the Boltz-2 confidence metrics, ipTM-min held the most discriminative performance (Figure 1) consistently outperforming other metrics when used as a single decision value for distinguishing native vs NCs, achieving 0.59 ROC-AUC for the non-crystalised part of the dataset and 0.61 ROC-AUC on the full dataset (Figure 2). The full dataset demonstrates inflated performance, so the rest of this manuscript focuses on the non-crystallized part of the dataset; follow-up analyses are provided provided in the supplementary section for the full dataset.

**Figure 1:**
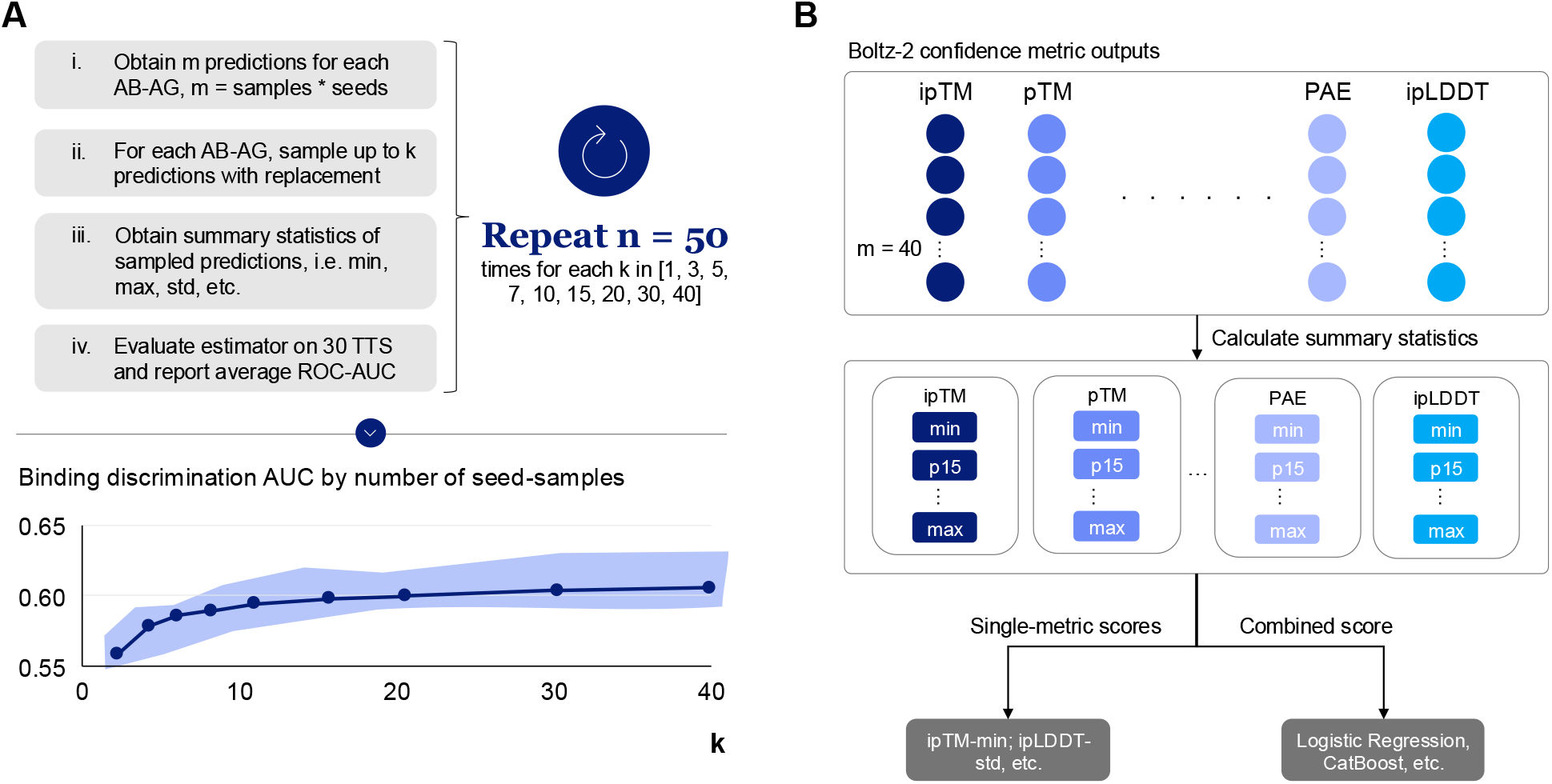
Overview of the unified experimental setup for evaluating discriminative power and estimating uncertainty of Boltz-2 metrics and classifiers with increasing numbers of seeds and samples. The left-hand side **(A)** covers the Monte-Carlo evaluation loop whilst the right-hand side **(B)** describes the estimators.

**Figure 2:**
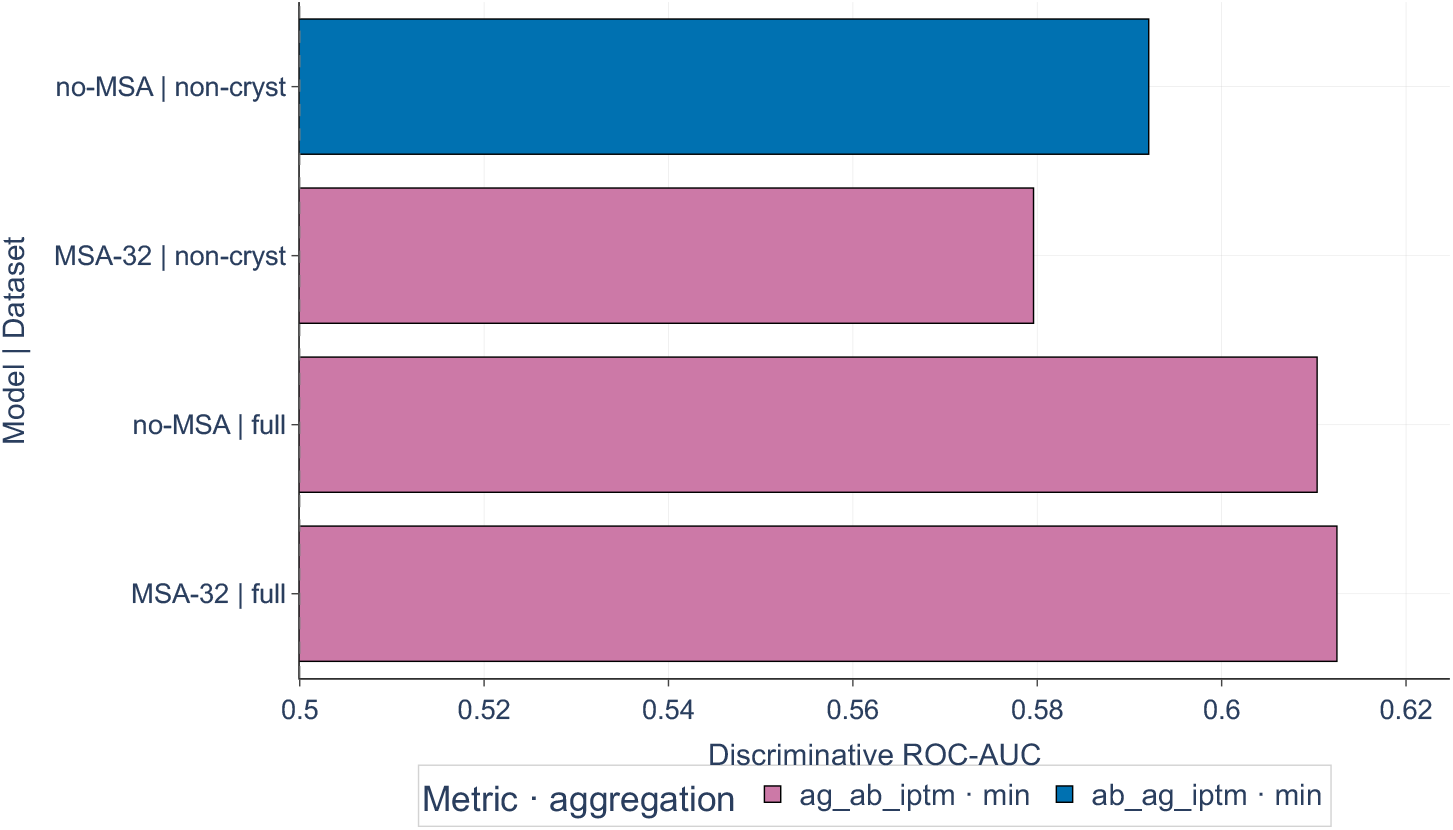
Baseline ROC-AUC performances using individual Boltz confidence metrics across different model settings (MSA-32 vs no-MSA) and across datasets (full vs non-crystalized structures only). The classification task is classifying native vs negative control antibody-antigen pairs.

### Multivariate classifier performance

Multiple classification models were evaluated, however, none consistently outperformed the simple ipTM-min baseline (SI Table 2), as the observed differences were small relative to the associated variance. Under Monte-Carlo evaluation, logistic regression demonstrated the best overall performance showing modest improvement over ipTM-min baseline (SI Figure 2). With the logistic regression and across classifiers, performance plateaued at approximately 0.6 ROC-AUC when 40 seed-samples were used, compared to an initial value of approximately 0.55 ROC-AUC with a single seed-sample (Figure 4). The single-metric baselines, Ab– Ag ipTM-min and Ag–Ab ipTM-min, remained difficult to outperform and constituted the strongest-performing approaches among models incorporating either MSA without structural information or structural information without MSA on the full dataset. These baselines exhibited lower uncertainty than the multivariate classifiers, as expected as they were not learning relationships from the training data.

Uncertainty bounds of approximately ±0.01 ROC-AUC were observed across all classifiers, arising from variability in the training data and differences in test set partitioning. All models consistently outperformed the lower region around 0.5 ROC-AUC observed under permuted labels (SI Figure 2), thereby confirming performance above chance level.

The superior performance of logistic regression under the Monte-Carlo evaluation scheme suggests that, in low-sample regimes and under variability induced by multi-seed-sample inference, only simple parametric models can be estimated well with the size of the dataset. Furthermore, a marked improvement in performance was observed as the number of seed-samples increased from one to twenty; however, only marginal gains were achieved beyond this point, with slight additional improvements up to approximately thirty seed-samples (Figure 4; SI Figure 2).

Given the limited performance gains beyond twenty seed-samples, comparative analyses of classifier performance were conducted at this level (SI Figure 3). At this stage, differences in the relative performance of estimators are more clearly discernible. Although the mean ROC-AUC values of several classifiers exceeded those of the baseline methods, the corresponding percentile ranges overlap, indicating limited statistical separation between approaches.

For the data without structures, performance obtained using MSA was consistently lower across the trajectory, although a similar overall trend was observed (Figure 4). In contrast, when considering the full dataset (SI Figure 2), the difference in performance between models with and without MSA was less pronounced. Furthermore, under these conditions, certain classifiers trained on outputs incorporating MSA achieved the highest performance at 40 seed-samples.

### Impact of MSA on performance

Hypothesis testing was conducted on the data at 20 seed-samples, corresponding to the point at which performance had reached a plateau. Model performance with and without MSA was compared across the different estimators (Figure 3).

**Figure 3:**
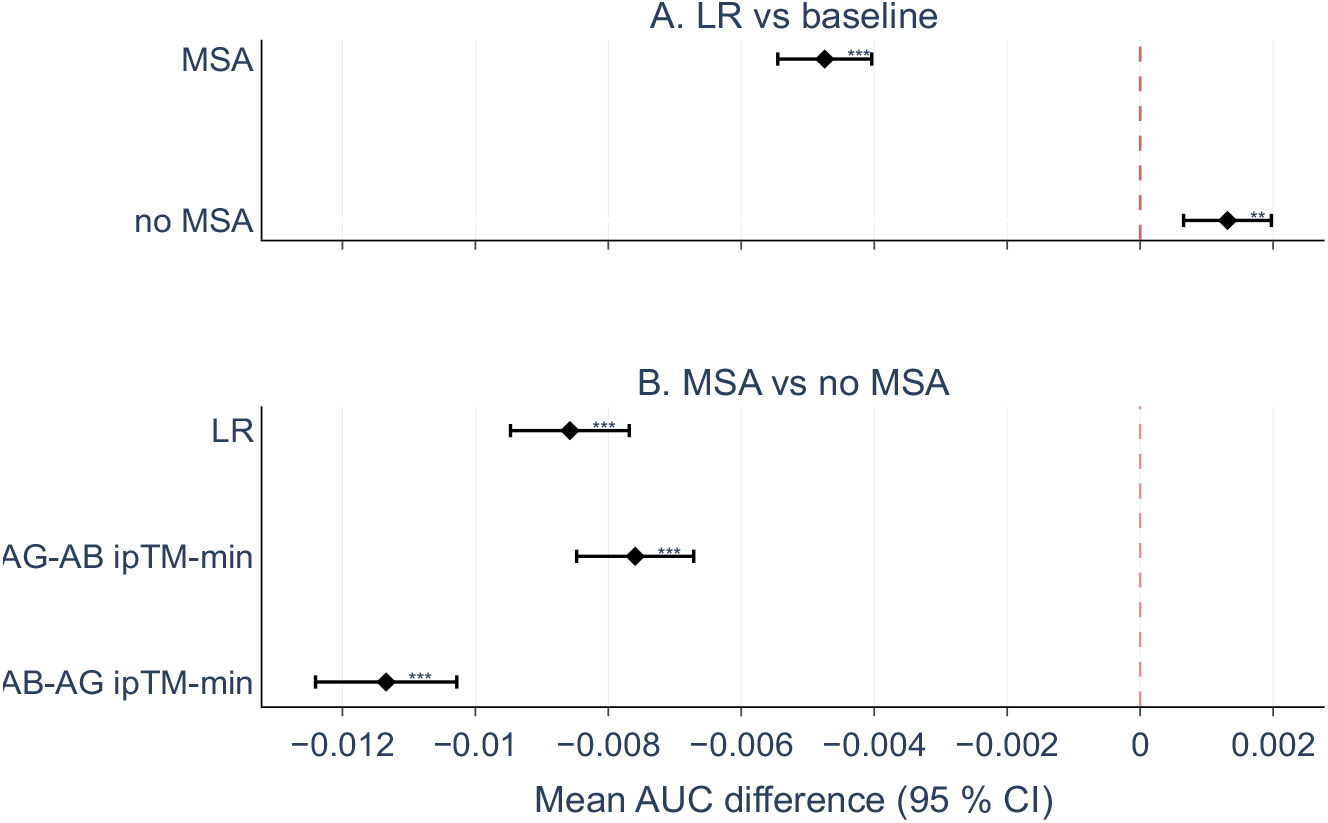
Results of testing discriminative performance over different model settings and classifier options. Hypothesis test used was a two-sided paired t test comparing mean AUC of the configuration at step 20, i.e. 20 seed-samples. P values were adjusted using Bonferroni correction within the step and across all comparisons. Significance levels are annotated using adjusted p values with levels *** if p < 0.001, ** if p < 0.01 and * if p < 0.5 else ns. For example, the top panel (A) compares the performance of mean ROC-AUC of LR model minus mean of ROC-AUC of baseline model with and without MSA.

**Figure 4:**
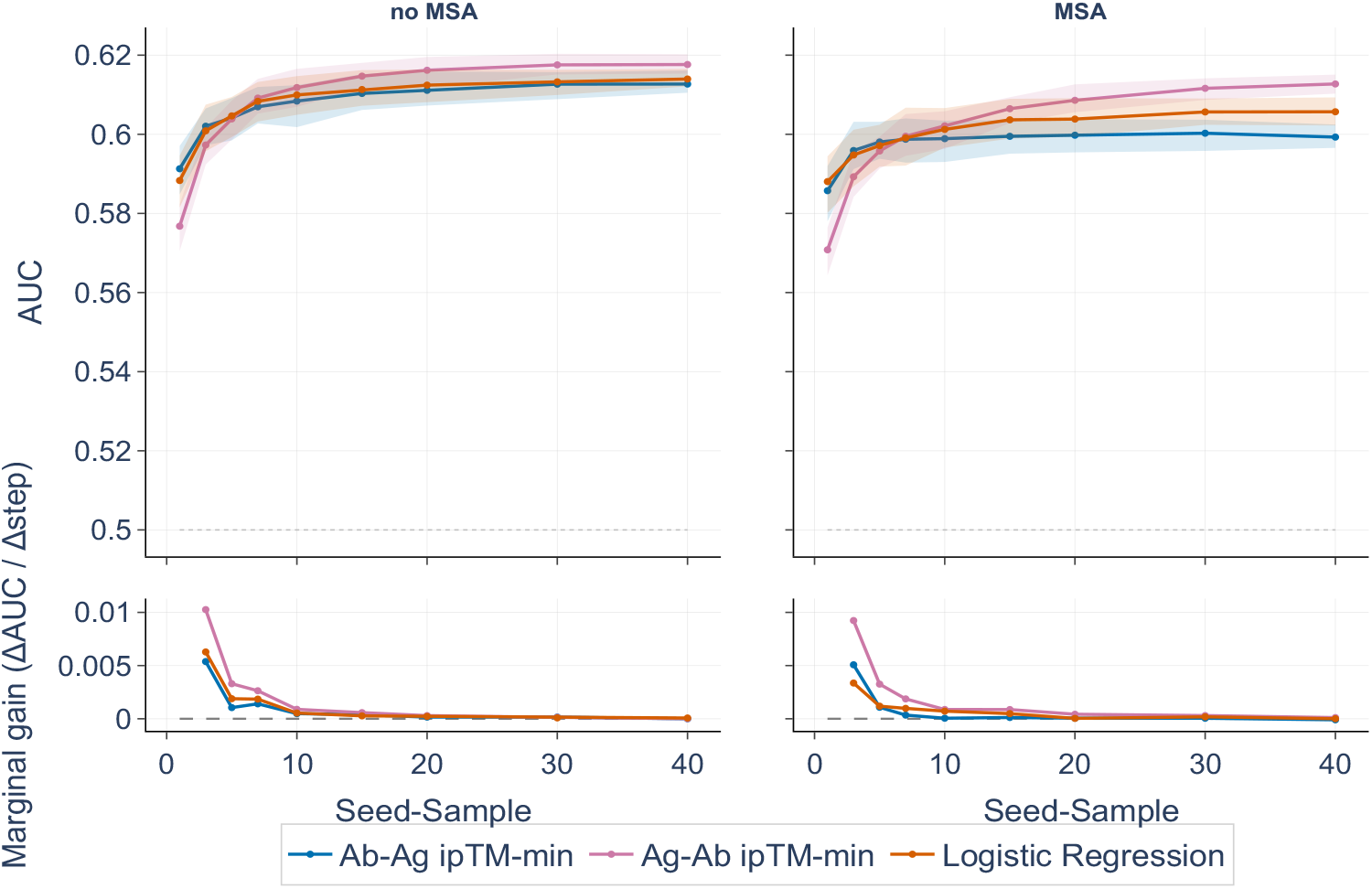
ROC-AUC performances of logistic regression vs baseline models using raw Boltz-2 confidence metrics across different model settings (MSA-32 vs no-MSA) and for increasing sampling depth. Uncertainty bounds are built using 5th and 9th percentiles across bootstrap samples. Marginal gain shows performance increase in the average ROC-AUC score between consecutive steps. The data used is without structures only and the classification task is classifying native vs negative control antibody-antigen pairs.

At 20 seed-samples, the inclusion of 32 MSAs was found to significantly reduce discriminatory performance (Figure 3). This effect was consistently observed across ipTM-based metrics as well as in the logistic regression model. The observed changes in ROC-AUC ranged from +0.0049 to +0.0342 across the conducted tests, while effect sizes, as measured by Cohen’s d, were approximately −1.08 to −10.87 (SI Figure 4). Furthermore, it is important to note that on the structure-excluded subset, inference time excluding MSAs was ∼8s faster on average.

### Comparison of logistic regression vs baselines

An additional comparison was performed between classifier-based approaches and baseline metrics. The logistic regression model was found to significantly outperform the top ipTM-min baseline under conditions without MSA; however, this advantage was not observed when MSA was included (Figure 3). The magnitude of improvement in ROC-AUC was less than 0.005, corresponding to an effect size of approximately Cohen’s d ≈ +1.19. This indicates a narrow and limited benefit, rather than a substantial or systematic improvement in modeling performance.

## Discussion

This work presents a systematic exploration of Boltz-2 inference settings in a novel benchmarking framework for AB– AG discrimination, including controlled evaluation across key dimensions such as sampling depth, model-derived confidence metrics (e.g., ipTM, pTM), MSA usage, and dataset composition. Across multiple Boltz-2 output metrics, including ipTM and aggregated confidence scores, we observed limited discriminative performance, with ROC-AUC values in the 0.55-0.62 range indicating only the weak discriminative performance despite being above chance. These results suggest that, under deployable conditions, Boltz-2 confidence metrics may provide limited utility for identifying true binders. Importantly, this conclusion is supported by a robust evaluation framework that explicitly accounts for variability across stochastic seeds, repeated train– test splits, and bootstrapped resampling of prediction sets, enabling stable estimation of performance across realistic screening scenarios. More broadly, this underscores the challenges of applying general-purpose structure prediction outputs to antibody discovery tasks and motivates further development of approaches optimized for binding discrimination.

These limitations can be understood in the context of fundamental properties of AB–AG interactions. Such interfaces exhibit substantial structural diversity, ranging from relatively flat epitopes to deep pockets engaged by highly flexible CDR loops, requiring accurate prediction of both loop conformations and complementary epitope surfaces. Unlike natural protein–protein interactions, therapeutic antibodies are engineered against targets that may lack meaningful co-evolutionary signals, which could reduce the utility of MSAs and aligning with the improved performance observed under no-MSA conditions.^28^ Additionally, current structure prediction models are trained to optimize structural accuracy rather than binding affinity, limiting their ability to distinguish biologically meaningful interactions from geometrically plausible but non-functional interfaces.^29^ These challenges are further compounded by biases in available structural data, where AB–AG complexes represent a small and non-uniform subset of protein–protein interactions.

Consistent with this interpretation, a recent benchmarking study of nanobody–antigen interactions similarly reports that internal confidence metrics such as ipTM fail to reliably distinguish cognate from non-cognate pairs, often assigning uniformly high scores across both classes.^17^ Notably, increased stochastic sampling improves structural refinement but does not enhance binding discrimination, suggesting that current confidence metrics are not calibrated to specificity. Our work extends these observations by systematically evaluating inference settings and introducing a screening-oriented framework that captures variability across seeds and configurations, while also controlling for confounding factors such as antigen length. Together, these results provide converging evidence across antibody formats that structure-derived confidence metrics alone are insufficient for reliable binding classification.

In practice, however, structure-derived confidence metrics are sometimes interpreted using threshold-based heuristics to classify interactions as binding or non-binding, but selecting an appropriate cutoff is challenging and highly dependent on dataset composition and class balance.^15,16,30^ In contrast, ROC-AUC provides a threshold-independent measure of discriminative performance, and the near-random values observed here indicate that no universal threshold yields consistent separation between binders and non-binders. As a result, even when a reasonable cutoff can be identified for a specific dataset, it is unlikely to generalize across screening scenarios.

Building on this, multivariate classifiers such as logistic regression can be used to combine multiple Boltz-2–derived features, effectively learning a weighted combination to distinguish native from non-cognate pairs. However, these approaches showed minimal improvement over individual metrics. This likely reflects a combination of low signal-to-noise ratio, label noise in the negative control set where some pairs may represent weak or uncharacterized binders, and a noisy, indirect relationship between structure-derived metrics and true binding affinity. Additionally, many of the most informative features are highly collinear, limiting the benefit of combining them, while biological and dataset-level heterogeneity—including variation in epitope accessibility, antigen flexibility, and unmodeled factors such as glycosylation or conformational state—further obscures consistent patterns. Together, these factors suggest that aggregating existing confidence metrics is insufficient for accurate classification without features explicitly linked to binding specificity.

To better contextualize these results, it is important to consider how the dataset and evaluation setup were constructed. Negative controls were constructed by randomly pairing antibodies with non-cognate antigens from the same pool, which controls for confounding factors such as antigen class and length but does not fully capture the complexity of true non-binding interactions. This approach may include both trivially non-interacting pairs and rare unrecognized binders and does not represent challenging negatives such as homologous or structurally similar antigens. As a result, the reported performance likely reflects an upper bound under simplified screening conditions. This limitation stems from the limited availability of experimentally validated non-binding AB–AG pairs in current training datasets, which is often due to the fact that non-binding interactions are less frequently reported or systematically characterized. It is worth noting that several companies have begun to develop antibody design platforms with lab-in-the-loop capabilities that will systematically generate richer training data. In addition, the analysis relied on a pretrained Boltz-2 model using sequence-based inputs with optional MSA information, without incorporating experimental constraints such as epitope mapping or affinity data. While this reflects realistic high-throughput settings, incorporating prior knowledge, such as restricting predictions to candidate epitope regions, may improve discrimination by reducing the search space and limiting spurious high-confidence interfaces.

Taken together, these findings suggest that meaningful improvements in antibody screening will require approaches explicitly optimized for binding discrimination rather than reliance on general structure-confidence metrics. This includes training objectives that incorporate binding specificity, development of curated negative datasets, and evaluation on application-relevant screening scenarios. Future work integrating experimentally validated datasets and more challenging negative controls will be critical for assessing performance under realistic conditions and advancing the utility of structure-based methods in antibody discovery.

## Methods

### Data Preparation and Modeling

Antibody sequences for 519 therapeutic monoclonal antibodies were retrieved from Thera-SabDab,^25^ a comprehensive database of therapeutic antibody structures and sequences. The dataset was filtered to include only monoclonal antibodies with a single protein target that could be modelled by Boltz-2; antibodies targeting whole viruses, small molecules, DNA, or lipids were excluded. Of these 519 antibodies, 143 had experimentally determined crystal structures available in the Protein Data Bank. Importantly, 129 of these structures were posted before the Boltz-2 training set cut-off date and therefore are likely to have been included in the Boltz-2 training dataset. For each antibody, both heavy and light chain sequences were obtained. Corresponding target antigen sequences were identified and retrieved from UniProt based on target identifiers annotated in Thera-SabDab, resulting in 225 unique target proteins across the dataset, whose crystallographic status remains unknown.^26^

Native AB–AG pairs were assembled using an automated sequence retrieval pipeline that programmatically extracted antigen sequences from UniProt based on annotated targets. A random negative control was generated using the same antigen pool by sampling a non-cognate antigen per antibody from the set of native targets, including repetitions.

### Boltz-2 model version used

All structure predictions were performed using Boltz-2 accessed through the NVIDIA Inference Microservices (NIM) release 1.4.0. The NIM implementation was selected for its computational efficiency, providing approximately 20% faster inference times compared to standard Boltz-2 deployment whilst maintaining performance,^31^ which is critical for large-scale screening applications involving hundreds of AB-AG pairs. Average inference time for no-MSA setup was around 24.5 seconds and 32.8 seconds for shallow MSA setup (32-MSA) per pair on NVIDIA H100s.

### Exploration of model parameters and sampling strategies

Boltz-2 predictions can vary based on several input settings that influence structure quality and diversity: seeds, samples, MSA, structural templates and user-defined constraints. The number of random seeds and diffusion samples per seed controls how extensively conformational space is explored with stochastic trajectories. MSAs provide evolutionary information by capturing coevolutionary constraints between interacting residues, potentially guiding the model toward biologically relevant conformations. Additionally, user-defined constraints such as distance restraints or contact predictions can explicitly guide structure generation.

This study focused on evaluating sampling strategies and MSA settings, as these are most critical for large-scale screening applications where computational efficiency, reproducibility, and applicability to novel AB-AG pairs are important.^19,32^

To examine variability introduced by random initialization, 8 random seeds and 5 diffusion samples per seed, collectively referred to as seed-samples, were generated for each AB-AG pair, yielding 40 total predictions per pair. This sampling strategy enabled quantification of structural variability across independent predictions, which enabled computation of statistics, including mean and standard deviation of confidence scores that provided discriminative signals for distinguishing native from negative control AB-AG pairs.^33^

MSA settings directly impact the evolutionary information available to guide predictions and have been shown to significantly affect model confidence metrics and prediction quality.^20^ Importantly, there can be a trade-off with the usage of MSAs as they can both guide and constrain a prediction. Although prior work has demonstrated MSA effects on structure prediction quality, the impact of MSAs on AB– AG binding discrimination remains unclear. Two factors complicate this question: First, diffusion-based models such as AlphaFold3 and Boltz-2 do not have EvoFormer modules and therefore do not intrinsically rely on co-evolutionary information in the same way as AlphaFold2. To evaluate MSA effects, two conditions were tested: (1) no MSA - structure prediction without evolutionary information and (2) shallow MSA - evolutionary context restricted to a depth of 32 sequences. In case 2, the MSA of the target sequence was provided as well to provide complete binding partner information. Template-based prediction was not explored in this study, as homologous AB–AG complex structures are often unavailable for novel therapeutic targets. Additionally, CDR/pocket constraints were not utilized as the precise epitope is often not known at inference.

### Seed variability analysis and evaluation approach

To evaluate the effect of stochastic sampling depth on native vs. negative control discrimination performance, a Monte Carlo simulation framework with Bootstrapping that models the variability arising from subsampling diffusion predictions (Figure 1) was implemented. For each AB–AG pair, 40 structural predictions across seed-samples were available and treated as stochastic samples of the diffusion process. To assess the impact of aggregating varying numbers of predictions, a sampling parameter was defined, representing the number of runs aggregated per AB–AG pair.

A total of 30 fixed train/test splits were generated using an 80/20 partition at the antibody level, ensuring that all data associated with a given antibody (both native and negative controls) were assigned exclusively to either training or testing. For each Monte Carlo iteration (n = 50 simulations) and for each value of *k* in 1, 3, 5, 7, 10, 15, 20, 30, 40 bootstrap resampling was performed by sampling *k* predictions with replacement from the available 40 runs for each AB– AG pair. Aggregated features (e.g., mean, standard deviation, and quantiles derived from confidence scores) were computed from the sampled runs.

For each of the 30 predefined splits and 50 simulations, different estimators were used to classify native vs random negative controls (see Estimators section for more details). As an additional negative control, class labels in the training data were randomly permuted within each split prior to model fitting, while evaluation was performed on the corresponding held-out test set. The ROC-AUC discrimination metric was computed for each simulation. Final reported performance corresponds to the mean of these 30 splits estimates, together with the empirical 5th–95th percentile range across the 50 Monte Carlo simulations; this range reflects variability under the resampling scheme rather than a formal confidence interval. To compare alternative model configurations (with and without MSA input) and alternative feature aggregation strategies (ipTM alone versus multivariate classifier score), simulations were paired by using identical subsampling indices across conditions. For each combination, paired performance differences (mean ROC-AUC) between configurations were computed using matched resampling across conditions. These paired differences were summarized across Monte Carlo replicates and assessed using paired t-tests (following the Central Limit Theorem). To control for multiple hypothesis testing, a Bonferroni correction was applied across the four hypotheses evaluated at each sampling depth. Effect sizes were quantified using the paired form of Cohen’s d.

### Metrics Analysis

The Boltz-2 model outputs a set of confidence metrics used to assess the prediction, many derived from the initial AlphaFold models.^9,18^ The primary outputs analysed in this study were the ipTM, pTM, pLDDT, PDE and interface predicted error (iPDE), and the model confidence score.

These metrics quantify model reliability at multiple structural scales. The pTM score reflects confidence in the global topology and domain organization of the predicted complex, whereas ipTM evaluates the predicted accuracy of inter-chain orientation and interface geometry. The pLDDT score provides per-residue confidence estimates describing local structural reliability. The PDE matrix captures expected pairwise positional uncertainty between residues. The confidence score represents an aggregate model-level ranking metric derived from internal structural confidence signals.

For each AB–AG pair, metric values were extracted for all generated samples across seeds and diffusion replicates. To account for stochastic variability, summary statistics were computed across corresponding sampled predictions: the median, mean, standard deviation, minimum, maximum value, 15th and 85th percentiles.

The resulting aggregated statistics were used as features in downstream classification and ranking analyses, both individually and in combination, to evaluate their discriminative capacity in native versus negative-control tasks.

### Estimators

#### Baselines

A set of Boltz-2 confidence metrics (pTM and ipTM, pLDDT and ipLDDT, pDE and ipDE, confidence score) were used as heuristic baselines for AB–target discrimination. Each metric was aggregated across the AB–AG, AG–AB or VH–VL interfaces, yielding multiple versions of each metric (e.g. AB– AG ipTM vs AG–AB ipTM).

Each baseline score was aggregated with a fixed set of summary operators: max, min, mean, median, standard deviation and percentiles at both 15 and 85. The resulting single-value scorer was then compared against the native/negative-control labels using ROC–AUC.

#### Stacked classifiers

A set of classifiers were trained on each of the train sets in the Monte Carlo simulation and taking aggregated Boltz-2 metrics as features. The models chosen were logistic regression, CatBoost, random forest, extremely randomized trees (Extra Trees) and linear SVM (to have coverage of linear and tree-based ensemble models). The hypothesis behind these stacked classifiers was that they could leverage multiple Boltz-2 metrics to provide improved discriminative performance than a single baseline metric.

### Fine-tuning and Feature Selection

Model tuning was performed outside of the Monte Carlo bootstrap using the same fixed train/test split with consistent grouped folds. Performance was evaluated as the mean ROC-AUC across folds.

First a randomized joint search was conducted over feature subsets and hyperparameters for logistic regression and CatBoost. Feature subsets were defined by sampling groups of related variables, retaining 25–100% of groups per trial. The best-performing CatBoost configuration was then used for SHAP-based recursive feature selection, applied independently per fold and aggregated to obtain a consensus feature set by choosing top 15 features. Final model selection was carried out via a grid search across multiple model families (logistic regression, linear SVM, random forest, Extra Trees, CatBoost, and baseline scorers) using the selected features. Models were trained and evaluated on each fold with fold-specific standardization to avoid data leakage. Performance was summarized as mean ± std across folds, with ROC-AUC as the primary metric. The selected features and tuned hyperparameters can be found in SI Table 1 and SI Table 3 respectively.

## Acknowledgements

We would like to thank Harmeet Atwal, Rasmus Hildebrandt, and Benedikt Dietz for their valuable contributions to shaping the problem statement and early guidance on the approach. We also thank Saira Qureshi and Aditi Gautam from NVIDIA for their support with the NIM implementation of Boltz-2.

