## Supplemental Information for "Benchmarking Boltz-2 for Screening of Therapeutic Antibody-Antigen Interactions"

### Supplementary File

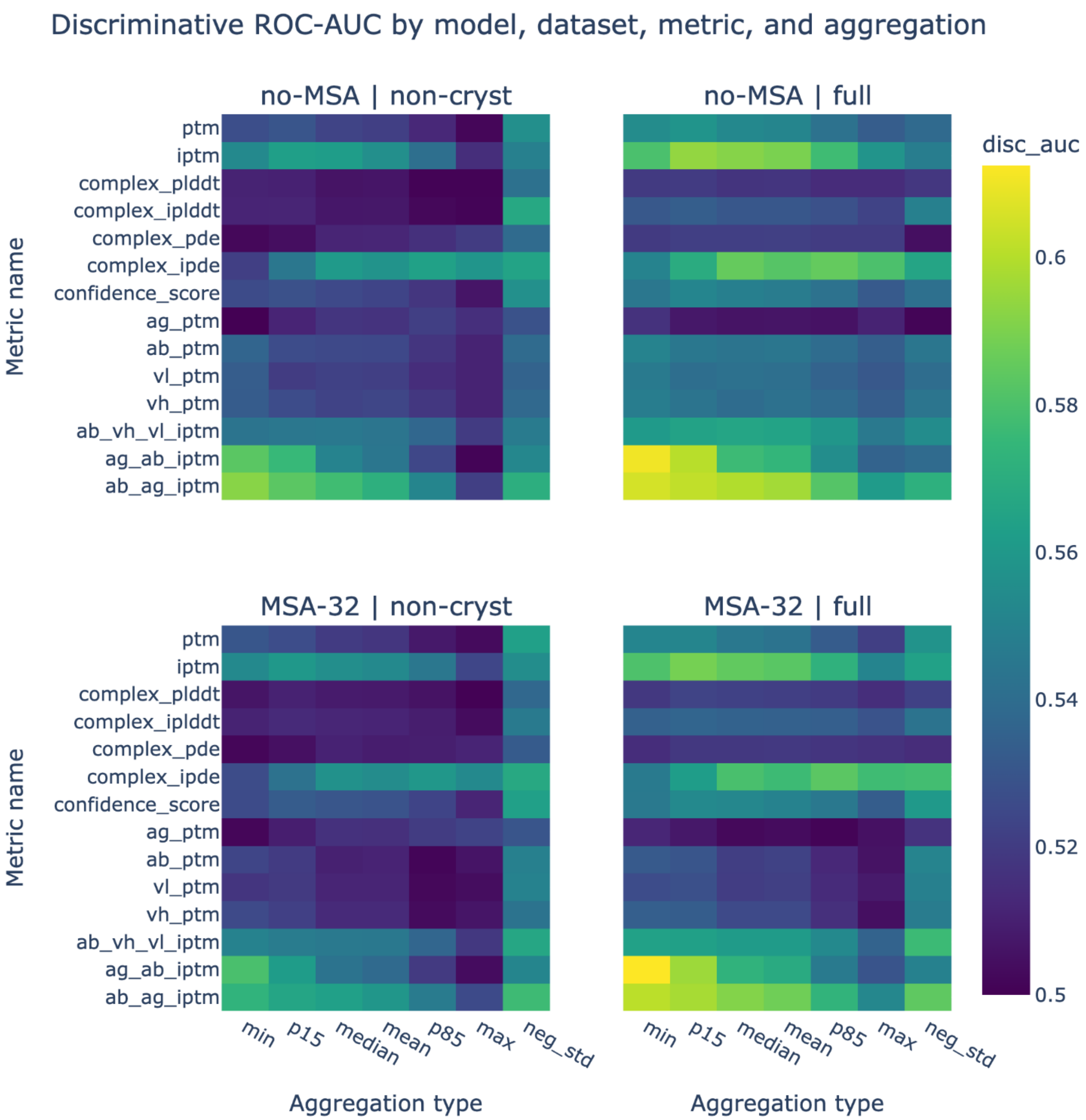


**Figure S1: ipTM aggregated across AB-AG and AG-AB interfaces using minimum across samples was the most discriminative metric.** Performance here is measured using median AUC-ROC across data from Boltz-2 predictions with and without MSA, with classification task of classifying native vs negative control antibody-antigen pairs.

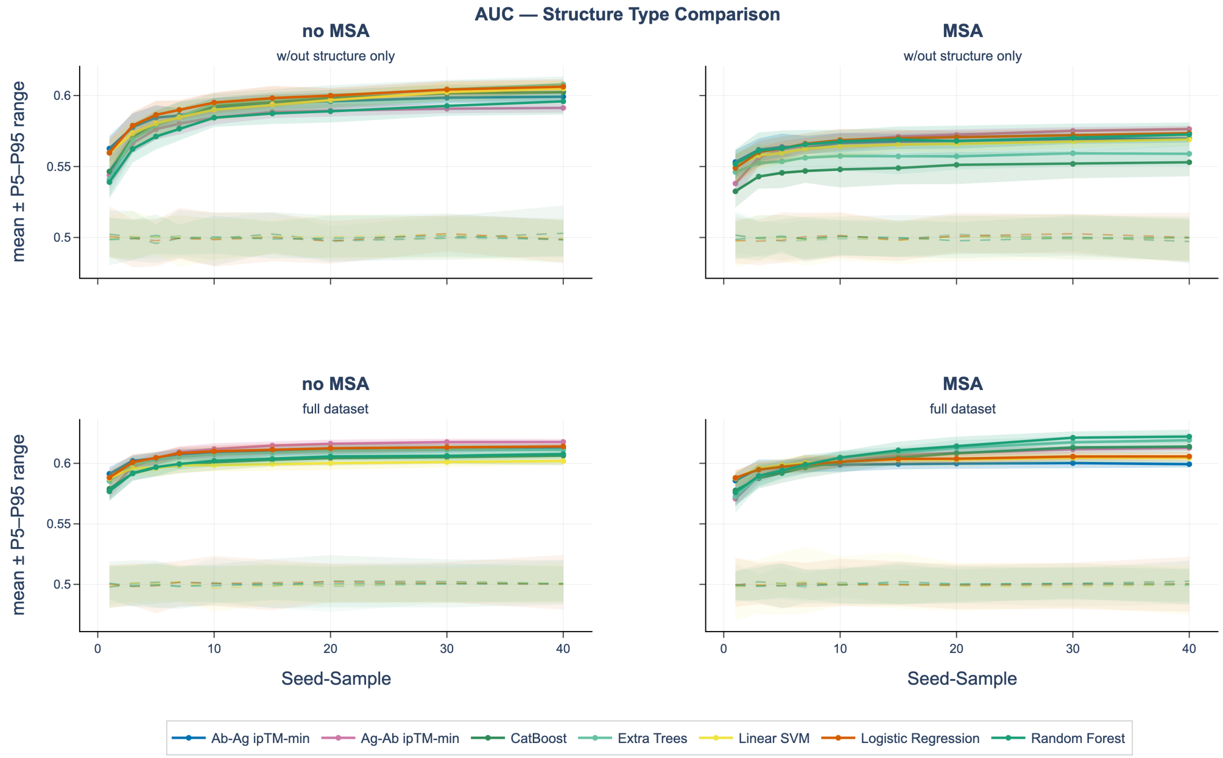


**Figure S2: ROC-AUC performances all classifiers vs baseline models using raw Boltz-2 confidence metrics across different model settings (MSA-32 vs no-MSA) and for increasing sampling depth.** Uncertainty bounds are built using 5th and 95th percentiles across bootstrap samples. The data used is without structures only and the classification task is classifying native vs negative control antibody-antigen pairs. The shaded region shows the AUC of classifiers trained on data with shuffled labels.

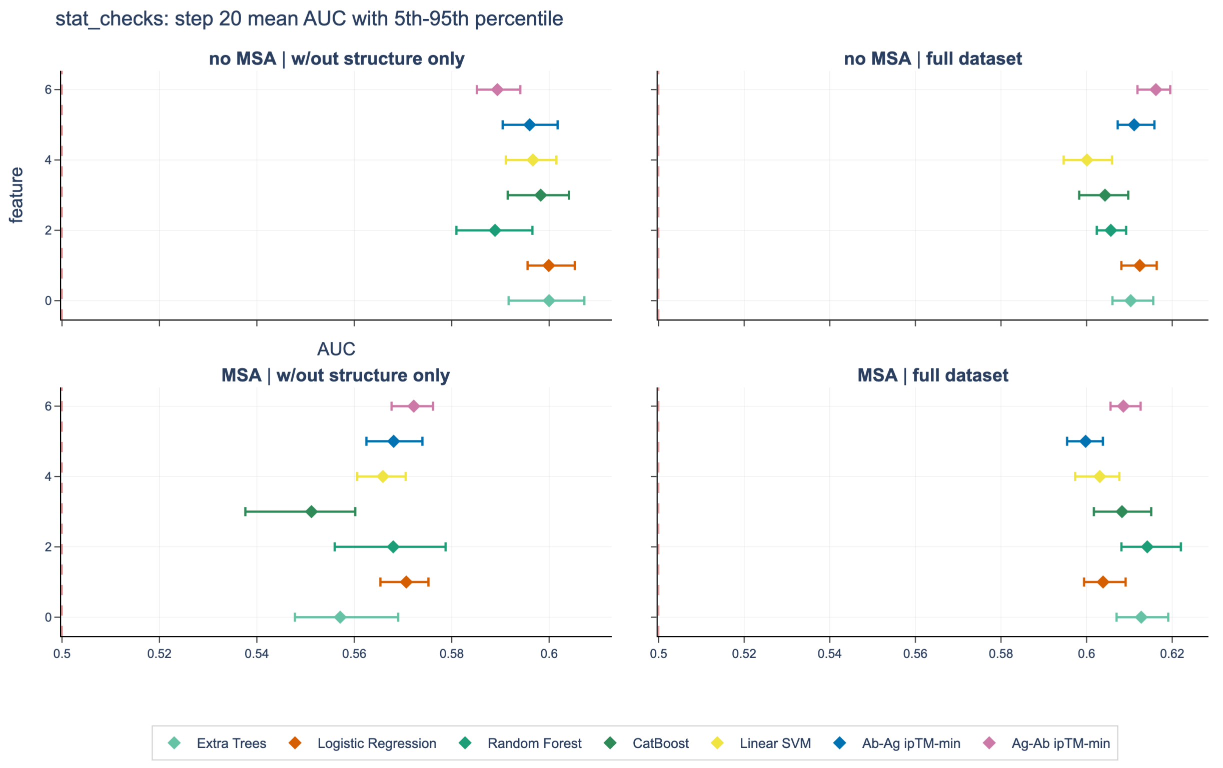


**Figure S3: Performances of classifiers and baselines on the 20 seed-samples step across MSA / no MSA model settings and data without structures and full data.** Mean AUC and $5^{th}$-$95^{th}$ percentiles across the samples are plotted.

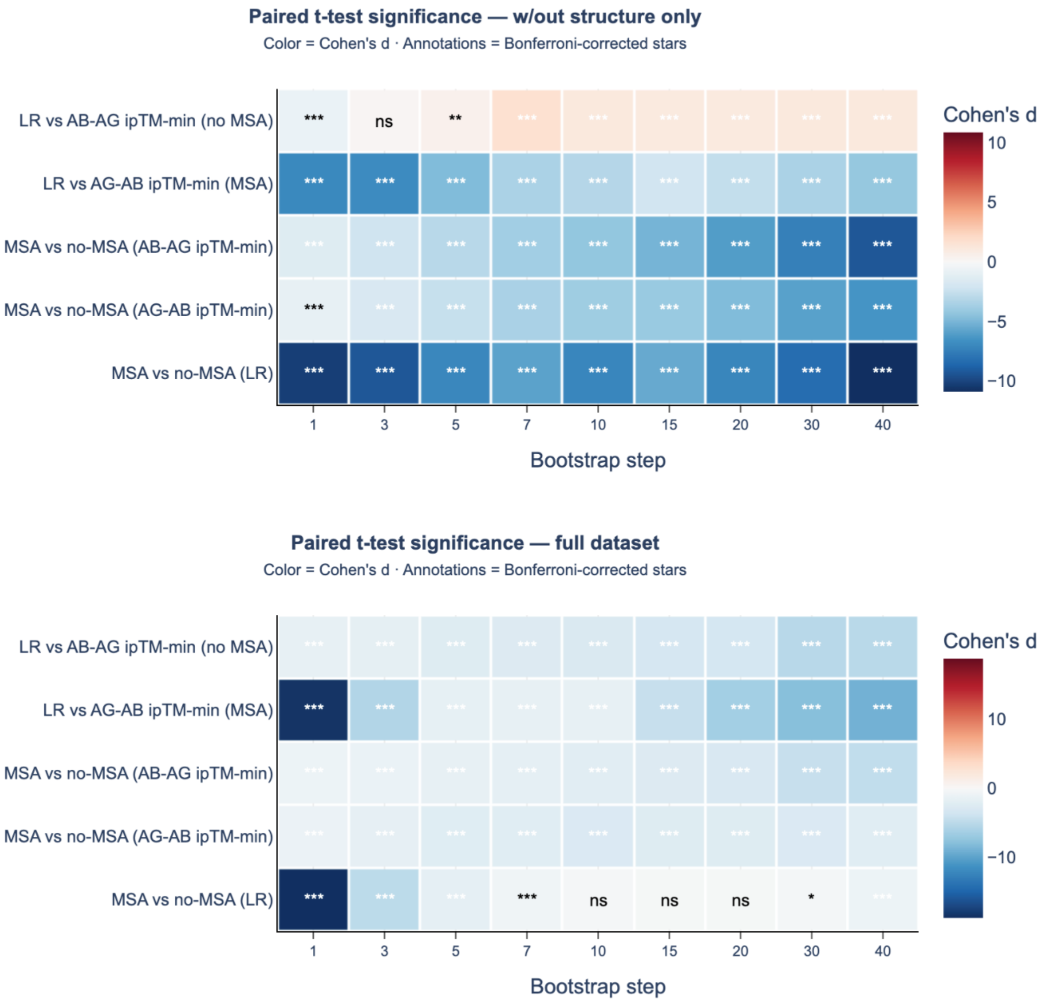


**Figure S4: Effect sizes (using Cohen's D) comparing performance of estimator scenarios.** Each test is a two-sided paired t-test comparing average AUC of the two estimators, and significance is adjusted using Bonferroni correction within a bootstrap step. Significance levels are annotated using adjusted p values with levels *** if $p < 0.001$, ** if $p < 0.01$ and * if $p < 0.05$ else ns.

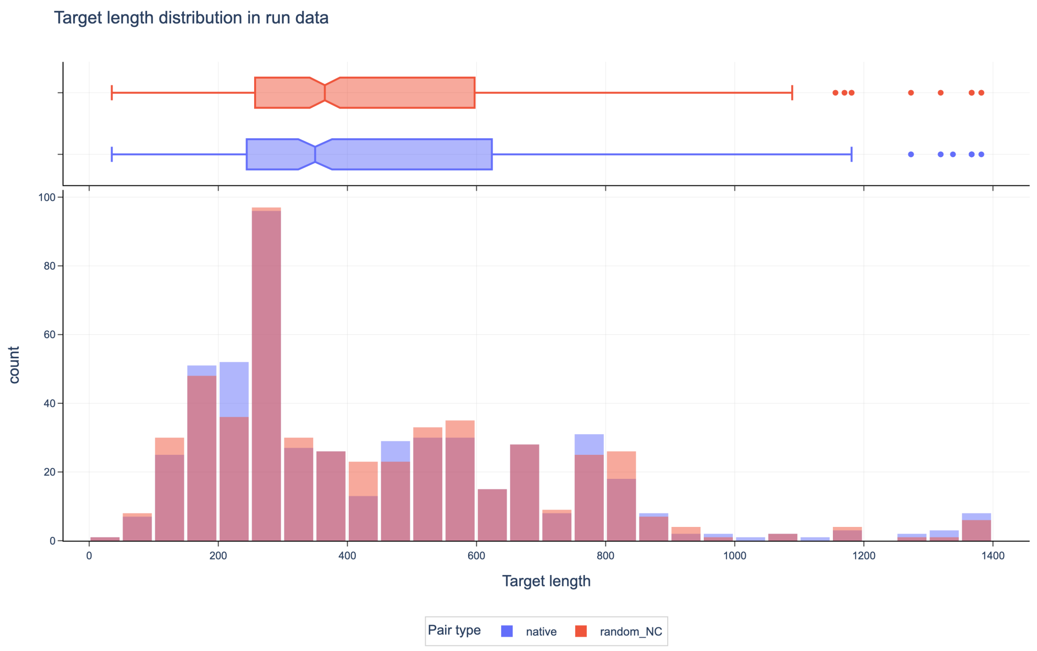


**Figure S5: The antigen length distributions for the two classes of interest, native and negative controls, are similar.** This is consistent with the previous finding that antigen length has negligible predictive value (ROC-AUC is 0.51).

| model_setting | no-MSA (non-cryst) | MSA-32 (non-cryst) | no-MSA (full) | MSA-32 (full) |
|---|---|---|---|---|
| ab_ag_iptm__min | 0.6001 ± 0.0450 | 0.5683 ± 0.0482 | 0.6131 ± 0.0312 | 0.5987 ± 0.0406 |
| ag_ab_iptm__min | 0.5909 ± 0.0349 | 0.5801 ± 0.0465 | 0.6177 ± 0.0343 | 0.6165 ± 0.0410 |
| catboost | 0.6172 ± 0.0413 | **0.5905** ± 0.0549 | **0.6213** ± 0.0343 | **0.6351** ± 0.0278 |
| extra_trees | **0.6193** ± 0.0416 | 0.5858 ± 0.0443 | 0.6184 ± 0.0255 | 0.6262 ± 0.0335 |
| linear_svm | 0.6171 ± 0.0324 | 0.5685 ± 0.0463 | 0.6077 ± 0.0320 | 0.6057 ± 0.0334 |
| logistic_regression | 0.6142 ± 0.0335 | 0.5732 ± 0.0487 | 0.6156 ± 0.0288 | 0.6070 ± 0.0336 |
| random_forest | 0.6136 ± 0.0473 | 0.5845 ± 0.0383 | 0.6144 ± 0.0302 | 0.6348 ± 0.0288 |

**Table S1: Best performing models from hyperparameter tuning.** The best model for each scenario is highlighted in bold. Chosen parameters were using in the Monte Carlo evaluation loop.

| Feature | MSA-full-data | MSA-no-structure | no-MSA-full-models | no-msa-no-structure |
|---|---|---|---|---|
| ab_ag_iptm__min | + | + | + | + |
| ab_ag_iptm__mean | + | + | | + |
| ag_ab_iptm__min | + | + | + | + |
| ab_ag_iptm__q0.15 | + | + | + | + |
| complex_ipde__std | + | + | + | |
| complex_iplddt__std | + | | | + |
| iptm__q0.15 | + | | | |
| complex_ipde__mean | + | | | |
| ag_ab_iptm__std | + | | | |
| ab_vh_vl_iptm__q0.15 | + | | | |
| ab_ag_iptm__q0.85 | + | | | |
| ab_vh_vl_iptm__std | + | | | |
| iptm__min | + | | | |
| vh_ptm__max | + | | | |
| ag_ab_iptm__mean | + | | | |
| vl_ptm__std | | + | + | + |
| vl_ptm__q0.85 | | + | + | + |
| complex_plddt__min | | + | + | + |
| complex_pde__std | | + | + | + |
| vl_ptm__q0.15 | | + | | + |
| vl_ptm__min | | + | + | + |
| complex_plddt__mean | | + | + | + |
| complex_plddt__max | | + | + | + |
| complex_pde__q0.85 | | + | | + |
| complex_pde__q0.15 | | + | | |
| vh_ptm__std | | + | | + |
| vl_ptm__mean | | + | | |
| ag_ab_iptm__q0.15 | | | + | |
| complex_plddt__q0.85 | | | + | |
| complex_ipde__q0.85 | | | + | + |
| complex_plddt__q0.15 | | | + | + |

Table S2: Selected features from hyperparameter tuning to be used across tuned classifiers in the Monte Carlo evaluation loop.

| Model / Feature | no-MSA \| non-cryst | MSA-32 \| non-cryst | no-MSA \| full | MSA-32 \| full |
|---|---|---|---|---|
| ab_ag_iptm__min | column: ab_ag_iptm__min | column: ab_ag_iptm__min | column: ab_ag_iptm__min | column: ab_ag_iptm__min |
| ag_ab_iptm__min | column: ag_ab_iptm__min | column: ag_ab_iptm__min | column: ag_ab_iptm__min | column: ag_ab_iptm__min |
| catboost | depth: 6; iterations: 100; l2_leaf_reg: 3; learning_rate: 0.01; rsm: 0.85; subsample: 1.0 | depth: 3; iterations: 3; l2_leaf_reg: 5; learning_rate: 0.01; rsm: 0.7; subsample: 0.7 | depth: 2; iterations: 10; l2_leaf_reg: 10; learning_rate: 0.01; rsm: 0.85; subsample: 1.0 | depth: 5; iterations: 50; l2_leaf_reg: 1; learning_rate: 0.1; rsm: 1.0; subsample: 1.0 |
| extra_trees | class_weight: balanced; max_depth: None; max_features: sqrt; min_samples_leaf: 2; min_samples_split: 5; n_estimators: 200 | class_weight: balanced; max_depth: 8; max_features: 0.5; min_samples_leaf: 10; min_samples_split: 2; n_estimators: 5 | class_weight: balanced; max_depth: 3; max_features: 0.3; min_samples_leaf: 2; min_samples_split: 10; n_estimators: 50 | class_weight: balanced; max_depth: 8; max_features: 0.5; min_samples_leaf: 1; min_samples_split: 5; n_estimators: 200 |
| linear_svm | C: 1.0; class_weight: balanced; loss: hinge | C: 0.01; class_weight: balanced; loss: hinge | C: 0.1; class_weight: balanced; loss: squared_hinge | C: 0.01; class_weight: balanced; loss: hinge |
| logistic_regression | C: 1.0; class_weight: balanced; l1_ratio: 1.0; solver: liblinear | C: 0.01; class_weight: balanced; l1_ratio: 0.0; solver: liblinear | C: 0.01; class_weight: balanced; l1_ratio: 0.0; solver: saga | C: 0.01; class_weight: balanced; l1_ratio: 0.0; solver: liblinear |
| random_forest | class_weight: balanced; max_depth: 12; max_features: log2; min_samples_leaf: 5; min_samples_split: 2; n_estimators: 200 | class_weight: balanced; max_depth: None; max_features: 0.3; min_samples_leaf: 5; min_samples_split: 5; n_estimators: 50 | class_weight: balanced; max_depth: 3; max_features: log2; min_samples_leaf: 2; min_samples_split: 10; n_estimators: 100 | class_weight: balanced; max_depth: 5; max_features: 0.3; min_samples_leaf: 20; min_samples_split: 10; n_estimators: 50 |

**Table S1: Information by model and data scenario of best model parameters following hyperparameter tuning. Chosen parameters were using in Monte Carlo evaluation loop.**

### Supplementary Note 1: Reproducibility information

All analysis was done in Python 3.13 environments. The Boltz-2 implementation was NVIDIA Inference Microservices (NIM) release 1.4.0. Models were run with scikit-learn 1.8.0 and catboost 1.2.8. Hypothesis testing was run with scipy 1.17.0. Sequence data was handled using biopython 1.8.
